# Shared genetic etiology between obsessive-compulsive disorder, obsessive-compulsive symptoms in the population, and insulin signaling

**DOI:** 10.1101/608034

**Authors:** Janita Bralten, Joanna Widomska, Ward De Witte, Dongmei Yu, Carol A. Mathews, Jeremiah M. Scharf, Jan Buitelaar, Jennifer Crosbie, Russell Schachar, Paul Arnold, Mathieu Lemire, Christie L. Burton, Barbara Franke, Geert Poelmans

## Abstract

**Objective:** Obsessive-compulsive symptoms (OCS) in the population have been linked to obsessive-compulsive disorder (OCD) in genetic and epidemiological studies. Insulin signaling has been implicated in OCD. We extend previous work by assessing genetic overlap between OCD, population-based OCS, and central nervous system (CNS) and peripheral insulin signaling.

**Methods:** We conducted genome-wide association studies (GWASs) in the population-based Philadelphia Neurodevelopmental Cohort (PNC, 650 children and adolescents) of the total OCS score and six OCS factors from an exploratory factor analysis of 22 questions. Subsequently, we performed polygenic risk score (PRS) analysis to assess shared genetic etiologies between clinical OCD (using GWAS data from the Psychiatric Genomics Consortium), the total OCS score and OCS factors. We then performed gene-set analyses with a set of OCD-linked genes centered around CNS insulin-regulated synaptic function and PRS analyses for five peripheral insulin signaling-related traits. For validation purposes, we explored data from the independent Spit for Science population cohort (5047 children and adolescents).

**Results:** In the PNC, we found a shared genetic etiology between OCD and ‘impairment’, ‘contamination/cleaning’ and ‘guilty taboo thoughts’. In the Spit for Science cohort, we were able to validate the finding for ‘contamination/cleaning’, and additionally observed genetic sharing between OCD and ‘symmetry/counting/ordering’. The CNS insulin-linked gene-set associated with ‘symmetry/counting/ordering’. We also identified genetic sharing between peripheral insulin signaling-related traits (type 2 diabetes and the blood levels of HbA1C, fasting insulin and 2 hour glucose) and OCD as well as certain OCS.

**Conclusions:** OCD, OCS in the population and insulin-related traits share genetic risk factors, indicating a common etiological mechanism underlying somatic and psychiatric disorders.

## Introduction

Obsessive-compulsive disorder (OCD) is a heterogeneous psychiatric condition characterized by persistent, intrusive thoughts and urges (obsessions) and repetitive, intentional behaviours (compulsions) (1). OCD affects 2-3% of the world’s population (2, 3). OCD is moderately heritable, with approximately 40% of the phenotypic variance explained by genetic factors, and a higher genetic load has been reported in childhood onset OCD (4, 5). The genetic architecture of OCD is complex, with multiple genetic variants of small effect size contributing to its etiology. This has hampered the identification and replication of genetic susceptibility factors. A meta-analysis of hypothesis-driven candidate gene association studies has implicated serotoninergic and catecholaminergic genes in OCD, while studies focusing on glutamatergic and neurotrophic genes have shown inconsistent results (6). Neither of the two independent genome-wide association studies (GWASs) of OCD (7, 8) nor a subsequent meta-analysis (2688 cases and 7037 controls) (9) yielded genome-wide significant findings, likely due to lack of power. However, the meta-analysis demonstrated that the polygenic signal from either sample predicted OCD status in the other sample, indicating the polygenic nature of the disorder (9).

The diagnosis of OCD is based solely on clinical symptoms, and no genetic or biological markers are available with sufficient specificity and accuracy to be clinically actionable (1). However, factor analyses of OCD symptoms have consistently identified specific OCD symptom clusters or dimensions, with the most reliable including contamination/cleaning, doubt/checking, symmetry/ordering, and unacceptable/taboo thoughts (10–13).

Obsessive-compulsive symptoms (OCS) are also present in the general population (14–16). Indeed, 21 to 38 % of individuals in the population endorse obsessions and/or compulsions, although only a small minority (2-3 %) meet the DSM-5 criteria for a clinical OCD diagnosis (17, 18). OCS are also heritable, with their heritability ranging from 30 to 77% (19, 20). In addition to contributing to overall OCS, genetic factors contribute to specific OCS dimensions, including contamination/cleaning (21–23) and checking/ordering (23, 24). Genetic overlap between clinical OCD and OCS in the population is suggested by the fact that polygenic risk scores (PRS) based on OCD GWAS data significantly predict OCS in two population-based samples of 6931 and 3982 individuals, respectively (19, 25).

In a study aiming to identify molecular mechanisms underlying OCD, we earlier performed integration of the top ranked-results from the existing GWASs. This resulted in a ‘molecular landscape’ that suggested the involvement of genes regulating postsynaptic dendritic spine formation and function through central nervous system (CNS) insulin-dependent signalling (26). Support for a role of dysregulated insulin signaling in OCD and OCS comes from studies showing increased OCS in men with type 1 diabetes (27) and from a study indicating that OCD patients have a higher risk of developing type 2 diabetes (T2D) (28). Furthermore, OCS were found to be positively correlated with blood levels of glycosylated hemoglobin (HbA1c), a diagnostic measure of T2D (29), and OCD patients had markedly higher levels of fasting glucose (a characteristic of T2D) (30).

In this paper, we aimed to assess the presence and extent of genetic overlap between OCD, OCS in the population, and insulin signaling, using the largest available data sets. Specifically, we parsed phenotypic heterogeneity using an exploratory factor analysis of OCS measured in a population cohort of children and adolescents. Subsequently, we investigated the presence of shared genetic etiologies between OCD and the total and factorized OCS. We then assessed genetic sharing between OCD, OCS, and insulin-related traits. Lastly, we validated and extended our findings in an independent population cohort of children and adolescents.

## Methods

### Sample, phenotypic, and genetic data

We studied OCS in the Philadelphia Neurodevelopmental Cohort (PNC) (31–34), which includes 8719 children and adolescents aged 8-21 years with neurobehavioral phenotypes and genome-wide genotyping data. Participants in the PNC provided written consent for genomic studies when they presented to the Children’s Hospital of Philadelphia health care network. OCS were assessed with GO-ASSESS, a computerized version of the Kiddie-Schedule for Affective Disorders and Schizophrenia (K-SADs) (35). For the current study, we selected 22 GO-ASSESS questions that corresponded to the diagnostic criteria for OCD (**Supplementary Table 1**). Participants were included if they answered the questions related to obsessions and/or compulsions. If those questions were all answered “no”, we allowed the questions on the consequences of obsessions and compulsions to be left blank, as no consequences are expected if no symptoms are present. The scores for each of the questions (0 for “no” and 1 for “yes”) were then summed to create a total OCS score (range 0-22). Genome-wide genotyping in the PNC cohort was performed in waves using six different genotyping platforms (details in **Supplementary Methods**). As a primary aim of our study was to assess the genetic overlap between OCD and OCS in the population, we only used phenotypic and genetic data from those PNC participants who answered positively on at least one of the questions related to the presence of obsessions and/or compulsions. This resulted in a final sample of 650 individuals for the subsequent factor and genome-wide association analyses.

### Factor analysis

First, using SPSS 23 (SPSS Technologies, Armonk, NY, USA), we determined the internal consistency (Cronbach’s α) of the 22 questions that constitute the total OCS scores in the 650 PNC participants. We then conducted a factor analysis of the scores on the 22 questions using Promax rotation to determine the number of factors that, when combined, explains the largest portion of the observed variance in the total OCS score. Specifically, we considered scree plots, eigenvalues >1 and the cumulative variance explained when selecting the number of factors and assigned questions to factors based on the highest absolute loading value.

### Genome-wide association analyses

Quality control filtering was applied to the genetic data to remove single nucleotide polymorphisms (SNPs) with low minor allele frequency (MAF) (<0.05), poor genotype call rate (<95%), and deviations from Hardy-Weinberg equilibrium (P<1×10^−6^). The imputation protocol used MaCH (36) for haplotype phasing and minimac (37) for imputation. Imputed SNPs with low imputation quality score (info<0.6) and low MAF (<0.05) were removed. If the total OCS score or the scores for the OCS factors fell within the limits of a normal distribution (i.e., a skewness and kurtosis between −1 and 1), we used a continuous trait design for the genome-wide association analysis. Otherwise, we used a pseudo case-control design, in which all individuals with a score of 0 for a factor were defined as ‘controls’ and compared against the ‘pseudo cases’, i.e. all individuals with a score of 1 or more for that factor. GWASs were carried out with mach2qtl (36) using the total OCS score and the scores for those factors that showed sufficient variation as phenotypes, with age and gender included as covariates. GWASs were performed separately for each genotyping platform and combined in an inverse-variance-weighted meta-analysis using METAL (38), accounting for genomic inflation.

### Shared genetic etiology analyses

#### OCD and OCS

First, we determined the level of shared genetic etiology between diagnosed OCD and OCS in the population. For this, we used the summary statistics from the meta-analysis of the two published GWASs of OCD (9) (data provided through the Psychiatric Genomics Consortium (PGC) for 2688 OCD cases and 7037 controls) as the ‘base’ sample for polygenic risk score (PRS)-based analyses in PRSice (39). The summary statistics from the GWASs of the different OCS in the PNC were used as the ‘target’ samples for the PRS-based analyses. For details see **Supplementary Methods**.

Multiple comparisons correction was done using the Benjamini-Hochberg false discovery rate (FDR) method (40, 41).

#### OCD, OCS and peripheral insulin signaling-related traits

To determine the level of genetic sharing between five peripheral insulin signaling-related traits and OCD as well as OCS, we conducted PRS-based analyses in PRSice (39), as described above. As base samples, we used summary statistics data from GWASs of the following peripheral insulin signaling-related traits: type 2 diabetes (T2D) and the blood levels of four T2D markers: HbA1c, fasting insulin, fasting glucose and glucose 2 hours after an oral glucose challenge (2hGlu) (details in **Supplementary Methods**). As target samples of the PRS-based analyses, we used the summary statistics from the OCD GWAS meta-analysis and the GWASs of the total OCS score and the scores for the OCS factors in the PNC. P-values of shared genetic etiology were corrected using the Benjamini-Hochberg FDR method.

#### Gene-set analyses

We first compiled a set of all genes encoding proteins within our molecular landscape of OCD (26) (see above). This resulted in a set of 51 autosomal genes for subsequent analyses. Using the GWAS results of the total OCS score and the scores for the OCS factors, gene-set analyses were then performed using the Multimarker Analysis of GenoMic Annotation (MAGMA) software (42), see **Supplementary Methods**. P-values were considered significant if they exceeded a Bonferroni-corrected threshold accounting for the number of phenotypes tested (P < 0.05/7 tests (total OCS score and six OCS factors)=0.00714). For significant gene-set associations, we looked at the individual gene-wide P-values and applied Bonferroni correction (P < 0.05/51 genes in the gene-set=0.00098).

#### Validation analyses in an independent population sample

In order to validate and possibly expand our findings, we performed PRS-based and gene-set analyses using data from GWASs of OCS in an independent population sample: the ‘Spit for Science’ project which includes 16,718 children and adolescents aged 6-17 years recruited from a local science museum (43). OCS were measured using the Toronto Obsessive-Compulsive Scale (TOCS), a validated 21-item parent-or self-report questionnaire (14). TOCS items are scored from −3 (far less often than others of the same age) to +3 (far more often than others of the same age). We first assessed which TOCS questions could be grouped into OCS factors similar to those calculated based on the PNC data. Two OCS factors (‘symmetry/counting/ordering’ and ‘contamination/cleaning’) were similar, see **Supplementary Table 2**. Genome-wide genotyping data for 5047 individuals of Caucasian descent entered the ‘continuous trait’ GWAS analysis for each factor. A description of genotyping, quality control and imputation can be found elsewhere (44) and GWAS details in the **Supplementary Methods**. Using summary statistics of the GWASs of the two TOCS OCS factors, we examined the shared genetic etiology between OCD and the TOCS OCS factors, and between the five peripheral insulin signaling-related traits and the TOCS OCS factors. Gene-set analyses between the set of 51 genes from the OCD landscape and the two TOCS OCS factors were also performed.

## Results

### Factor analysis

The internal consistency between the scores on the 22 OCS questions from the PNC was satisfactory (Cronbach’s α=0.69). **Supplementary Figure 1A** shows the total score distribution (mean=6.4, s.d.=3.35). Factor analysis revealed an eight factors solution as the best-fitting model, explaining 58.6% of the variance in the total score. We named these eight OCS factors ‘impairment’, ‘symmetry/counting/ordering’, ‘contamination/cleaning’, ‘aggressive taboo thoughts’, ‘repetition’, ‘guilty taboo thoughts’, ‘distress’, and ‘religious taboo thoughts’ (**Table 1**; factor score distributions in **Supplementary Figure 1B**).

**Table 1.**
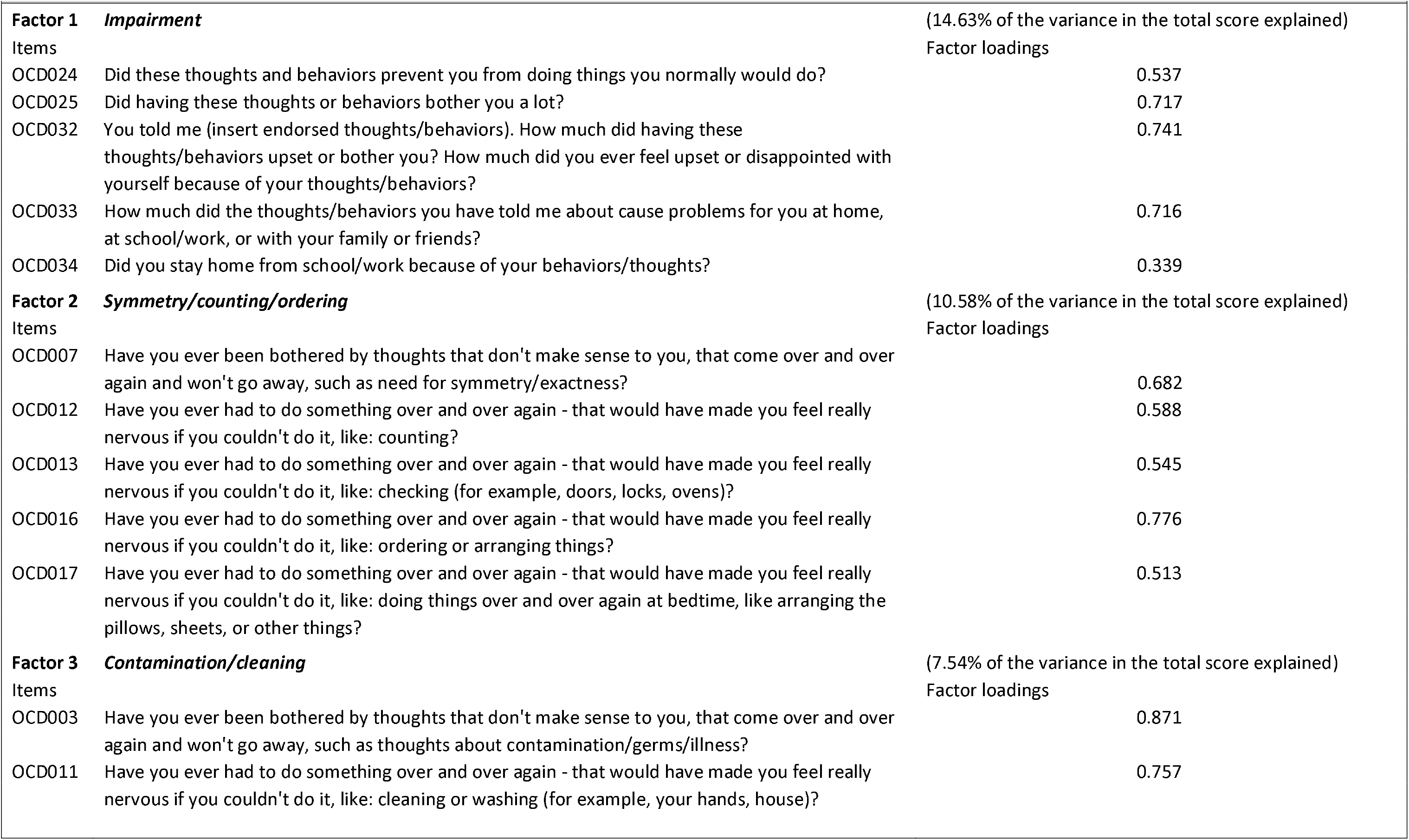
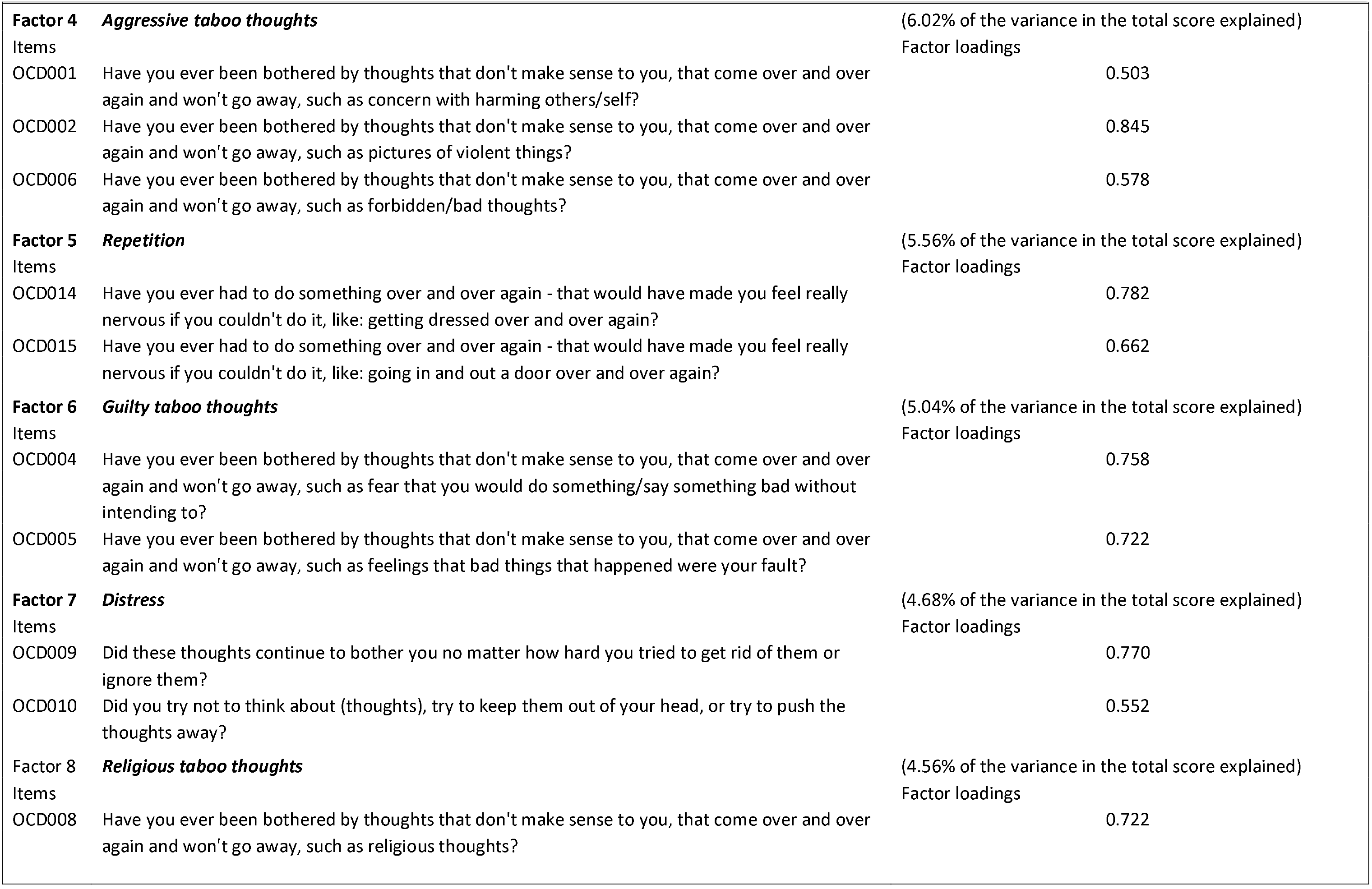
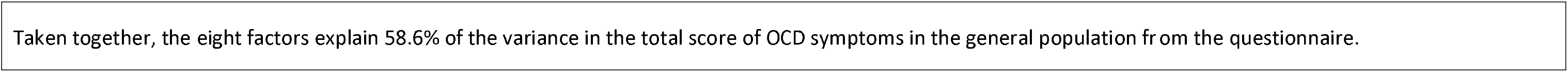
Item content of and loadings on the eight factors that constitute the best fitting model to explain the variance in the total score of the 22 items from the questionnaire of obsessive-compulsive symptoms that was completed by 650 participants from the PNC cohort.

### Genome-wide association analyses

Based on the distributions of the scores, we used a continuous trait design for the GWASs of the total OCS score and the factors ‘impairment’, ‘symmetry/counting/ordering’, and ‘guilty taboo thoughts’. A pseudo case-control design was used for the factors ‘contamination/cleaning’, ‘aggressive taboo thoughts’, and ‘distress’. The distribution of the scores on the OCS factors ‘repetition’ and ‘religious taboo thoughts’ showed too little variation to be taken forward (**Supplementary Figure 1B**).

### Shared genetic etiology analyses

#### OCD and OCS

We found statistically significant evidence for a shared genetic etiology between diagnosed OCD and three population-based OCS factors: ‘impairment’ (variance explained or R^2^ = 1.05%; FDR-adjusted P = 1.66E-02), ‘contamination/cleaning’ (R^2^ = 3.51%; FDR-adjusted P = 4.92E-06), and ‘guilty taboo thoughts’ (R^2^ = 4.89%; FDR-adjusted P = 1.41E-07) (**Supplementary Figure 2** and **Table 2**).

**Table 2.** PRS-based results for shared genetic etiology between OCD and the total OCS score as well as the scores for six OCS factors

|  | P<sub>T</sub> | P-value | R<sup>2</sup> | nSNPs |
| --- | --- | --- | --- | --- |
| Total OCS score | 0.05 | 4.79E-01 | 0.03% | 48168 |
| **Impairment** | **0.2** | **1.66E-02** | **1.05%** | **154269** |
| Symmetry/counting/ordering | 0.001 | 4.35E-01 | 0.05% | 1530 |
| **Contamination/cleaning** | **0.4** | **4.92E-06** | **3.51%** | **264566** |
| Aggressive taboo thoughts | 0.4 | 3.49E-01 | 0.13% | 264585 |
| **Guilty taboo thoughts** | **0.2** | **1.41E-07** | **4.89%** | **154267** |
| Distress | 0.1 | 1.55E-01 | 0.40% | 86806 |

<sup>#</sup> Shown in this table are the best SNP P-value thresholds (P<sub>T</sub>) for the PRSice analyses between OCD - 'base' sample - and the total OCS score and six OCS factors -'target' samples, their Benjamini-Hochberg adjusted P-values (P-value) for shared genetic etiology, the variance explained in the target sample phenotypes (R<sup>2</sup>), and the number of SNPs (nSNPs). Significant findings are indicated in bold.

#### OCD, OCS, and peripheral insulin signaling-related traits

We found statistically significant evidence for a shared genetic etiology between T2D and the total OCS score (R^2^ = 1.26%; FDR-adjusted P = 2.12E-02), ‘symmetry/counting/ordering’ (R^2^ = 3.98%; FDR-adjusted P = 3.23E-05), and ‘aggressive taboo thoughts’ (R^2^ = 3.62%; FDR-adjusted P = 3.54E-05) (**Supplementary Figure 3A** and **Table 3**). Blood HbA1c levels showed genetic sharing with total OCS score (R^2^ = 1.19%; FDR-adjusted P = 2.69E-02) and ‘aggressive taboo thoughts’ (R^2^ = 2.18%; FDR-adjusted P = 1.32E-03). Fasting insulin levels showed genetic sharing with OCD (R^2^ = 0.26%; FDR-adjusted P = 3.23E-05) and for fasting glucose levels, we did not find evidence of genetic sharing. Lastly, we observed genetic sharing between 2hGlu levels and OCD (R^2^ = 0.14%; FDR-adjusted P = 2.29E-03) as well as ‘guilty taboo thoughts (R^2^ = 1.12%; FDR-adjusted P = 2.96E-02) (**Supplementary Figure 3B-E** and **Table 3**).

**Table 3.** PRS-based results for shared genetic etiology between five peripheral insulin-signaling-related traits and OCD and OCS

| 'base' sample | 'target' sample | P<sub>T</sub> | P-value | R<sup>2</sup> | nSNPs |
| --- | --- | --- | --- | --- | --- |
| Type 2 Diabetes | OCD | 0.1 | 1.70E-01 | 0.03% | 138297 |
|  | **Total OCS score** | **0.5** | **2.12E-02** | **1.26%** | **322098** |
|  | Impairment | 0.5 | 4.47E-01 | 0.02% | 322098 |
|  | **Symmetry/counting/ordering** | **0.5** | **3.23E-05** | **3.98%** | **322098** |
|  | Contamination/cleaning | 0.5 | 2.38E-01 | 0.30% | 322043 |
|  | **Aggressive taboo thoughts** | **0.4** | **3.54E-05** | **3.62%** | **275379** |
|  | Guilty taboo thoughts | 0.05 | 4.03E-01 | 0.11% | 54161 |
|  | Distress | 0.5 | 2.94E-01 | 0.23% | 322098 |
| HbA1c | OCD | 0.001 | 4.03E-01 | 0.01% | 1152 |
|  | **Total OCS score** | **0.2** | **2.69E-02** | **1.19%** | **77169** |
|  | Impairment | 0.2 | 5.14E-02 | 0.95% | 77169 |
|  | Symmetry/counting/ordering | 0.4 | 4.03E-01 | 0.08% | 130416 |
|  | Contamination/cleaning | 0.001 | 1.15E-01 | 0.65% | 1126 |
|  | **Aggressive taboo thoughts** | **0.05** | **1.32E-03** | **2.18%** | **24321** |
|  | Guilty taboo thoughts | 0.1 | 4.03E-01 | 0.09% | 43920 |
|  | Distress | 0.5 | 7.84E-02 | 0.80% | 152062 |
| Fasting Insulin | **OCD** | **0.2** | **3.23E-05** | **0.26%** | **12558** |
|  | Total OCS score | 0.1 | 3.23E-01 | 0.19% | 7353 |
|  | Impairment | 0.001 | 4.03E-01 | 0.12% | 342 |
|  | Symmetry/counting/ordering | 0.4 | 4.46E-01 | 0.03% | 20910 |
|  | Contamination/cleaning | 0.1 | 4.03E-01 | 0.11% | 7353 |
|  | Aggressive taboo thoughts | 0.4 | 4.03E-01 | 0.11% | 20909 |
|  | Guilty taboo thoughts | 0.001 | 3.10E-01 | 0.21% | 342 |
|  | Distress | 0.1 | 1.61E-01 | 0.49% | 7353 |
| Fasting Glucose | OCD | 0.4 | 2.94E-01 | 0.02% | 21585 |
|  | Total OCS score | 0.3 | 4.03E-01 | 0.10% | 17010 |
|  | Impairment | 0.05 | 1.93E-01 | 0.40% | 4794 |
|  | Symmetry/counting/ordering | 0.4 | 1.14E-01 | 0.66% | 21114 |
|  | Contamination/cleaning | 0.001 | 8.13E-02 | 0.78% | 518 |
|  | Aggressive taboo thoughts | 0.05 | 1.84E-01 | 0.43% | 4794 |
|  | Guilty taboo thoughts | 0.5 | 2.38E-01 | 0.30% | 25109 |
|  | Distress | 0.4 | 2.49E-01 | 0.29% | 21114 |
| 2h Glucose | **OCD** | **0.5** | **2.29E-03** | **0.14%** | **24147** |
|  | Total OCS score | 0.2 | 4.03E-01 | 0.11% | 10703 |
|  | Impairment | 0.2 | 4.06E-01 | 0.07% | 10703 |
|  | Symmetry/counting/ordering | 0.4 | 2.70E-01 | 0.26% | 19482 |
|  | Contamination/cleaning | 0.05 | 5.97E-02 | 0.88% | 3348 |
|  | Aggressive taboo thoughts | 0.001 | 2.35E-01 | 0.31% | 210 |
|  | **Guilty taboo thoughts** | **0.1** | **2.96E-02** | **1.12%** | **5932** |
|  | Distress | 0.2 | 2.25E-01 | 0.33% | 10703 |

<sup>#</sup> Shown in this table are the best SNP P-value thresholds (P<sub>T</sub>) for the PRSice analyses between five peripheral insulin signalling-related traits - 'base' samples, and OCD as well as OCS factors - 'target' samples, their Benjamini-Hochberg adjusted P-values (P-value), the variance explained (R<sup>2</sup>) in the target sample phenotypes, and the number of SNPs (nSNPs). Significant findings are indicated in bold.

#### Gene-set analyses

MAGMA-based gene-set analysis for the CNS insulin signalling genes extracted from our earlier-defined OCD landscape containing 33,329 SNPs (effective number of SNPs after adjusting for LD structure=2,189) revealed a significant association with ‘symmetry/counting/ordering’ (P=0.0038). Within the significant gene-set, none of the individual genes showed gene-wide association (**Supplementary Table 3**). No significant associations were found with total OCS score or the five other OCS factors.

#### Validation analyses in an independent population sample

Two OCS factors were similar between the PNC and Spit for Science cohort, i.e. ‘symmetry/counting/ordering’ and ‘contamination/cleaning’ (**Supplementary Table 2** and **Supplementary Figure 4A-B**). Using summary statistics of the GWASs of these two factors, we found that diagnosed OCD shows genetic sharing with ‘symmetry/counting/ordering _TOCS_’ (R2 = 0.49%; FDR-adjusted P = 2.42E-05) and ‘contamination/cleaning _TOCS_’ (R2 = 0.23%; FDR-adjusted P = 4.07E-03), with the latter providing a validation of our finding in the PNC data.

We observed a shared genetic etiology between T2D and ‘contamination/cleaning _TOCS_’ (R2 = 0.28%; FDR-adjusted P = 1.59E-03) (**Supplementary Table 4** and **Supplementary Figure 5A-C**). Gene-set analysis for the OCD landscape genes in the two OCS _TOCS_ factors revealed no significant associations.

All results from the PRS-based analyses are summarized in **Table 4**.

**Table 4.**
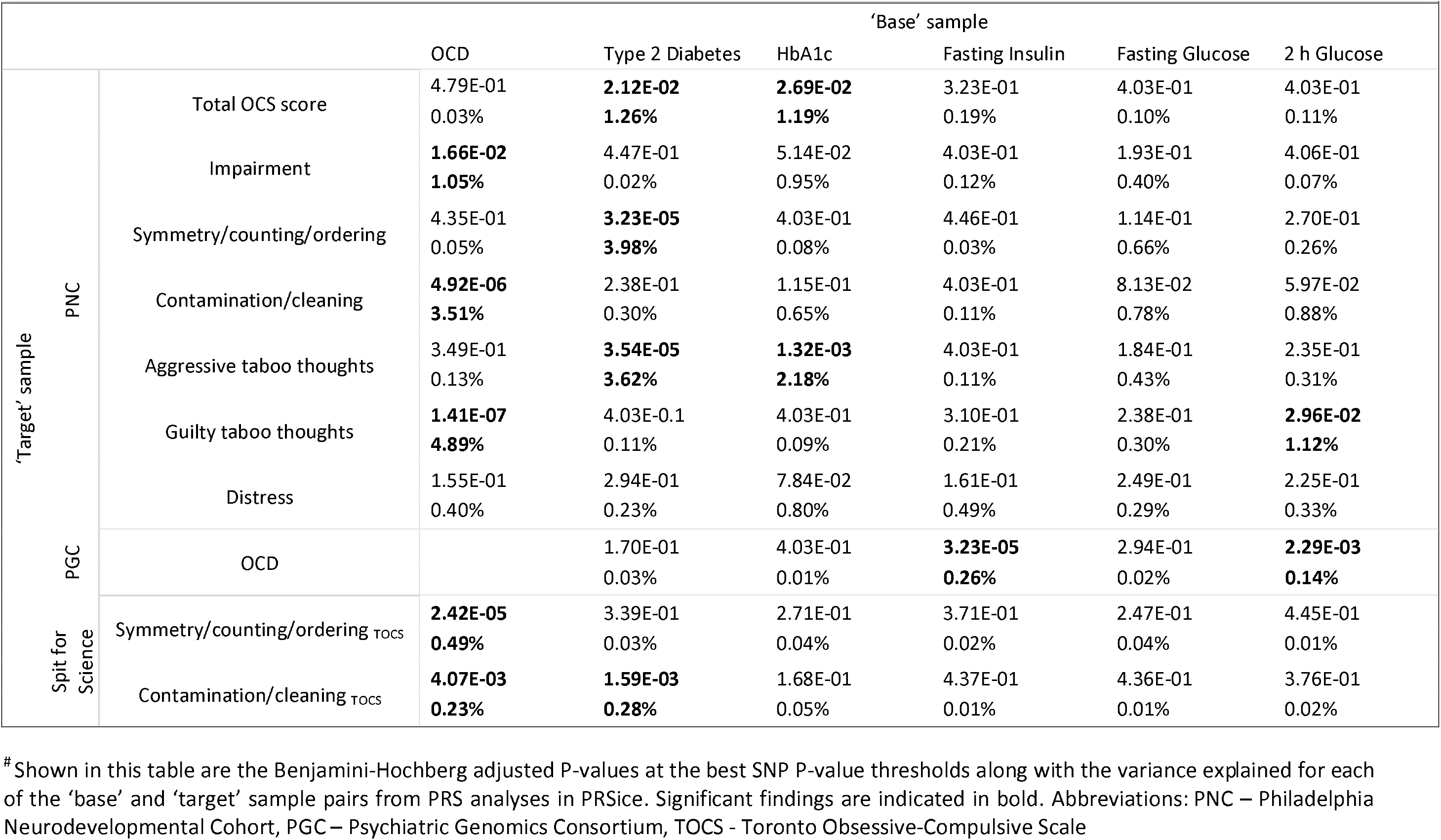
Summary of results from PRS-based analyses

## Discussion

In this study, we extended previous work by assessing genetic overlap between OCD, OCS in the population, and CNS and peripheral insulin signaling. While previous studies (19, 25) have yielded a shared genetic etiology between OCD and the total population-based OCS score, our analyses using phenotypic and genetic data of 650 children and adolescents from the population (PNC cohort) found genetic sharing between OCD and the OCS factors ‘impairment’, ‘contamination/cleaning’, and ‘guilty taboo thoughts’. We validated the finding for ‘contamination/cleaning’ in the larger Spit for Science cohort (n=5047). In this cohort, we also expanded our results by showing genetic sharing between OCD and ‘symmetry/counting/ordering’. Our findings are in keeping with the literature suggesting (at least partial) genetic overlap between OCD and population-based OCS (19, 21–23). Further studies are needed to dissect the phenotypic and genetic heterogeneity of OCD and population-based OCS. OCD and OCS have been linked to altered CNS and peripheral insulin signaling. We found significant association between the set of 51 OCD genes centered around CNS insulin-regulated synaptic function and ‘symmetry/counting/ordering’. As for peripheral insulin signaling, we found genetic sharing between T2D and - based on the PNC data - the total OCS score, ‘symmetry/counting/ordering’ and ‘aggressive taboo thoughts’, and - in the Spit for Science cohort - ‘contamination/cleaning’. For three out of the four T2D blood markers (HbA1c, fasting insulin and 2hGlu), we identified shared genetic etiologies with OCD and specific OCS factors. These findings provide support for ‘dysregulated’ peripheral insulin signaling as a biological process contributing to both OCD and population-based OCS. Further evidence for a role of (altered) peripheral insulin signaling in OCD etiology is suggested by the fact that selective serotonin reuptake inhibitors (SSRIs), the first-line pharmacological treatment for OCD, positively affect diabetic parameters when used to treat depressive symptoms in T2D (i.e. decreasing HbA1c levels and insulin requirement, and increasing insulin sensitivity) (45). Interestingly, SSRIs are particularly effective for treating harm-related obsessions, which are a part of ‘aggressive taboo thoughts’ (46). This is in line with our finding of genetic sharing between T2D as well as HbA1c levels and ‘aggressive taboo thoughts’. In addition, a recent study demonstrated that bilateral deep brain stimulation (DBS), a safe and effective treatment option for pharmaco-resistant OCD, not only reduced OCD symptoms but also decreased fasting insulin levels in the blood of both OCD patients with T2D and non-diabetic OCD patients (47). Moreover, insulin in the CNS - either entering from the periphery by crossing the blood brain barrier (48) or synthesized in the CNS (49) - has important non-metabolic functions, including modulating synaptic plasticity (50) and learning and memory (51, 52).

Although it is not clear yet what the relative contributions are of dysregulated peripheral and CNS insulin signaling to OCD and OCS, we recently demonstrated that compulsivity observed in Tallyho (TH) mice, a rodent model of T2D, is potentially linked to disturbances in insulin signaling. TH mice both displayed compulsive behaviour and increased glucose levels in their dorsomedial striatum, which could be due to decreased action of peripheral and/or CNS insulin, and the glucose levels correlated with compulsivity (53).

The current results should be viewed in light of some strengths and limitations. A strength is that we used quantitative symptom scores collected through questionnaires in the general population, which has enabled us to generate OCS phenotypes that we could then perform GWASs on. Using samples selected from the community may also reduce selection bias, which can occur when patient samples are analysed (e.g. individuals suffering from several comorbid disorders are more likely to present for clinical care) (54). A limitation of the current study is that the sample size of the GWASs is too small to discover new single genetic variant associations. However, this sample size was large enough to provide proof of concept for genetic sharing between OCD, OCS in the population, and insulin signaling. The fact that we were able to validate part of our results in an independent cohort adds credibility to our findings. Another limitation may be that the proportions of the variance in the target phenotypes being explained by the base phenotypes are quite small. However, these ‘variances explained’ are in fact (much) higher than those found in similar analyses, e.g. the PRS derived from a GWAS of OCD explained (only) 0.20% of the variance in OCS in a population sample (19). Moreover, as the variance explained is dependent on the size of the ‘base sample’ for the generation of the PRS (55), the observed variances explained with the still relatively small meta-GWAS of OCD as base sample may be underestimated.

In conclusion, we identified a shared genetic etiology between OCD, OCS in the population, and both CNS and peripheral insulin signaling. Our results imply that altered insulin signaling is not only relevant for somatic disorders but is also involved in the etiology of psychiatric disorders and related symptoms in the population, especially OCD and OCS. Further studies are needed to disentangle the contributions of peripheral and CNS insulin production and signaling to these disorders and symptoms.

## Acknowledgements

The Philadelphia Neurodevelopment Cohort (PNC) sample is a publicly available data set. Support for the collection of the data sets was provided by the National Institute of Mental Health (Grant No. RC2MH089983 to Raquel Gur, M.D., Ph.D., and Grant No. RC2MH089924 to Hakon Hakonarson, M.D., Ph.D.). All participants were recruited through the Center for Applied Genomics at Children’s Hospital of Philadelphia. The Spit for Science cohort was funded by the Canadian Institutes of Health Research (PDA: MOP-106573; RJS: MOP–93696), with additional funding related to this project provided by the Alberta Innovates Translational Health Chair in Child and Youth Mental Health (PDA). The research leading to these results also received funding from the European Community’s Seventh Framework People Programme (FP7-PEOPLE-2012-ITN) under grant agreement no. 316978 (TS-EUROTRAIN), the European Community’s Horizon 2020 research and innovation programme under grant agreements no. 728018 (Eat2BeNICE) and no. 667302 (CoCA), from the Innovative Medicines Initiative (IMI) 2 Joint Undertaking (H2020/EFPIA) under grant agreements no. 115916 (PRISM), no. 115300 (EU-AIMS), and no. 777394 (AIMS-2-TRIALS), as well as from the Netherlands Organization for Scientific Research (NWO) based on a Vici personal grant (grant number 016-130-669 to BF) and a pilot grant from the Dutch National Research Agenda for the NeuroLabNL route project (grant number 400 17 602).

