## Supplementary Methods for "Shared genetic etiology between obsessive-compulsive disorder, obsessive-compulsive symptoms in the population, and insulin signaling"

**Sample, phenotypic and genetic data**

We studied OCS in the Philadelphia Neurodevelopmental Cohort (PNC) (1-4), which includes 8719 children and adolescents aged 8-21 years with neurobehavioral phenotypes and genome-wide genotyping data. Participants in the PNC provided written consent for genomic studies when they presented to the Children’s Hospital of Philadelphia health care network. Notably, participants from Philadelphia Neurodevelopmental Cohort (PNC) were not recruited from psychiatric clinics, and the sample is not enriched for individuals who seek psychiatric help.

Genome-wide genotyping in the PNC has been performed in waves using six different genotyping platforms. Genotypes are available through the NIMH Database of Genotypes and Phenotypes (dbGaP), study accession ID phs000607.v1.p1. ([https://www.ncbi.nlm.nih.gov/projects/gap/cgi-bin/study.cgi?study_id=phs000607.v1.p1&phv= 194292&phd=&pha=&pht=3445&phvf=28&phdf=&phaf=&phtf=&dssp=1&consent=&temp=1](https://www.ncbi.nlm.nih.gov/projects/gap/cgi-bin/study.cgi?study_id=phs000607.v1.p1&phv=194292&phd=&pha=&pht=3445&phvf=28&phdf=&phaf=&phtf=&dssp=1&consent=&temp=1)). We assessed the phenotypic distributions per platform, and for this study used the two genetic platforms with the largest number of individuals with phenotypic data, HumanOmniExpress-12v1.0 (OmniExpress) and Human610_Quadv1_B (610Quad) (n=1989 and n=1257, respectively). Hence, genome-wide genotyping data were available for 3246 individuals who also completed the 22 OCD questions. Ancestry was addressed by only selecting individuals of self-reported European descent and by using multidimensional scaling (MDS) analysis. Ancestry outliers (i.e., of non-European descent) were excluded based on visual inspection of the first two principal components, leaving 3041 individuals for analyses (n=1860 from 610Quad and n=1181 from OmniExpress). The final sample of 650 individuals for the subsequent factor and genome-wide association analyses included 418 individuals from 610Quad and 232 from OmniExpress.

**Shared genetic etiology analyses**

Before generating polygenic risk scores with PRSice (5), clumping was performed using PLINK (6) to select independent index SNPs for each linkage disequilibrium (LD) block in the genome. Based on the significance level of the SNPs in the base sample, the index SNPs were selected and form clumps of all other SNPs that are within 500 kb and are in LD (r^2^>0.25).

PRSice was then used to generate polygenic risk scores for OCD that are the sum of genome-wide SNPs associated with OCD weighted by their effect sizes estimated from the PGC OCD meta-GWAS and only including SNPs that exceed seven broad P-value thresholds. The seven thresholds that were used are 0.001, 0.05, 0.1, 0.2, 0.3, 0.4, and 0.5. PRSice calculated P-values of shared genetic etiology between the base OCD phenotype and the target phenotypes (total score of OCS and scores for the OCS factors) for each of the seven P-value thresholds (P_T_). Subsequently, the calculated P-values for genetic sharing were aggregated and corrected for multiple comparisons using the false discovery rate (FDR) method (7, 8). To calculate the FDR, we applied the Benjamini-Hochberg method implemented through mne.stats.fdr_correction function in python.

To determine the level of genetic sharing between five peripheral insulin signaling-related traits and OCD as well as OCS, we also conducted PRS-based analyses in PRSice (5). As base samples, we used summary statistics data from GWASs of the following peripheral insulin signaling-related traits: type 2 diabetes (T2D) (GWAS of 26,676 cases and 132,532 controls) (9) as well as the blood levels of four T2D markers: HbA1c (GWAS of 123,665 general population subjects; increased HbA1c levels are a diagnostic measure of T2D) (10), fasting insulin (GWAS of 108,557 general population subjects; increasing fasting insulin levels lead to insulin resistance, the pathological hallmark of T2D) (11), fasting glucose (GWAS of 133,010 general population subjects; increased fasting glucose levels (hyperglycemia) are a characteristic of T2D) (11), and glucose 2 hours after an oral glucose challenge (2-hour glucose or 2hGlu) (GWAS of 42,854 general population subjects; 2hGlu, a clinical measure of glucose tolerance used in the diagnosis of T2D) (11).

**Gene-set analyses**

For gene-set analyses, we first compiled a set of all genes encoding proteins within our molecular landscape of OCD that is involved in regulating postsynaptic dendritic spine formation and function through CNS insulin-dependent signaling (12).

This resulted in a set of 52 unique genes but as our genome-wide association analysis only included autosomal SNPs, we had to omit one gene located on the X chromosome (*HTR2C*), leaving a final set of 51 genes for subsequent analysis. Gene-set analysis was then performed using the Multimarker Analysis of GenoMic Annotation (MAGMA) software (13). We selected all SNPs located within each of the 51 genes after which all single SNP P-values were transformed into a gene test-statistic by taking the mean of the χ2 statistic among the SNPs in each gene. Gene association tests were performed taking LD between SNPs into account using the 1000 Genomes Project European sample as a reference to estimate the LD between SNPs within the genes (<http://ctglab.nl/software/MAGMA/ref_data/g1000_ceu.zip>). Gene-wide P-values were converted to z-values reﬂecting the strength of the association each gene has with each of the OCS factors. Subsequently, we tested whether the 51 genes in the OCD landscape are jointly associated with the total score of OCS and the scores for the OCS factors. We used an intercept-only linear regression model, including a subvector corresponding to the genes in the gene-set. To test whether the association was different from the association of the genes outside the gene-set, taking into account the polygenic nature of our traits, the regression model was expanded to include all annotated genes in the genome outside the gene-set. To account for the potentially confounding factors of gene size and gene density, both gene size and gene density as well as their logarithms were included as covariates in the gene-set analysis.

**Validation analyses in an independent population sample**

To conduct the GWASs of the two OCS factors derived from the Spit for Science data, a linear regression model was used, adjusting for age, sex, respondent (questionnaire filled by parent or child), genotyping platform (Illumina HumanCoreExome-12 (v1-0) bead-chip or the HumanOmni1-Quad (v1.0) bead-chip), and the first 6 principal components from MDS.

The significance between the SNP imputed allele dosage and the response was calculated from a Wald test. Only SNPs with minor allele frequency > 0.01 and imputation quality (allelic R2) > 0.60 were tested.
