## Supplementary Table 1 for "Shared genetic etiology between obsessive-compulsive disorder, obsessive-compulsive symptoms in the population, and insulin signaling"

| **Supplementary Table 1.** Customized questionnaire of obsessive-compulsive symptoms in the general population. The 22 questions were from GO-ASSESS, a computerized version of the Kiddie-Schedule for Affective Disorders and Schizophrenia (K-SADs). | |
| --- | --- |
| **Nr** | **Questions** |
| OCD001 | Have you ever been bothered by thoughts that don't make sense to you, that come over and over again and won't go away, such as concern with harming others/self? |
| OCD002 | Have you ever been bothered by thoughts that don't make sense to you, that come over and over again and won't go away, such as pictures of violent things? |
| OCD003 | Have you ever been bothered by thoughts that don't make sense to you, that come over and over again and won't go away, such as thoughts about contamination/germs/illness? |
| OCD004 | Have you ever been bothered by thoughts that don't make sense to you, that come over and over again and won't go away, such as fear that you would do something/say something bad without intending to? |
| OCD005 | Have you ever been bothered by thoughts that don't make sense to you, that come over and over again and won't go away, such as feelings that bad things that happened were your fault? |
| OCD006 | Have you ever been bothered by thoughts that don't make sense to you, that come over and over again and won't go away, such as forbidden/bad thoughts? |
| OCD007 | Have you ever been bothered by thoughts that don't make sense to you, that come over and over again and won't go away, such as need for symmetry/exactness? |
| OCD008 | Have you ever been bothered by thoughts that don't make sense to you, that come over and over again and won't go away, such as religious thoughts? |
| OCD009 | Did these thoughts continue to bother you no matter how hard you tried to get rid of them or ignore them? |
| OCD010 | Did you try not to think about (thoughts), try to keep them out of your head, or try to push the thoughts away? |
| OCD011 | Have you ever had to do something over and over again - that would have made you feel really nervous if you couldn't do it, like: cleaning or washing (for example, your hands, house)? |
| OCD012 | Have you ever had to do something over and over again - that would have made you feel really nervous if you couldn't do it, like: counting? |
| OCD013 | Have you ever had to do something over and over again - that would have made you feel really nervous if you couldn't do it, like: checking (for example, doors, locks, ovens)? |
| OCD014 | Have you ever had to do something over and over again - that would have made you feel really nervous if you couldn't do it, like: getting dressed over and over again? |
| OCD015 | Have you ever had to do something over and over again - that would have made you feel really nervous if you couldn't do it, like: going in and out a door over and over again? |
| OCD016 | Have you ever had to do something over and over again - that would have made you feel really nervous if you couldn't do it, like: ordering or arranging things? |
| OCD017 | Have you ever had to do something over and over again - that would have made you feel really nervous if you couldn't do it, like: doing things over and over again at bedtime, like arranging the pillows, sheets, or other things? |
| OCD024 | Did these thoughts and behaviors prevent you from doing things you normally would do? |
| OCD025 | Did having these thoughts or behaviors bother you a lot? |
| OCD032 | You told me (insert endorsed thoughts/behaviors). How much did having these thoughts/behaviors upset or bother you? How much did you ever feel upset or disappointed with yourself because of your thoughts/behaviors? |
| OCD033 | How much did the thoughts/behaviors you have told me about cause problems for you at home, at school/work, or with your family or friends? |
| OCD034 | Did you stay home from school/work because of your behaviors/thoughts? |

**Explanatory Note**:

(1) The answers to questions OCD032 and OCD033 were re-categorized from a 10-point scale into binary responses (i.e., with scores from 0 to 4 converted into a score of 0 for "no" and scores from 5 to 10 converted into a score of 1 for "yes") for compatibility with the other 20 questions for which there were only two possible answers ("no"=score of 0, "yes"=score of 1).

(2) For the data selection, if the questions related to obsessions (OCD001-OCD008) and/or compulsions (OCD011-OCD17) were completed and all those questions were answered “no”, we allowed the questions on the consequences of obsessions and compulsions (OCD009, OCD010, OCD024, OCD025, OCD032, OCD033, OCD034) to be left blank, as no consequences are expected if no symptoms are present. The scores for each of the questions were then summed to create a total OCS score (range 0-22).
