## Supplementary Table 2 for "Shared genetic etiology between obsessive-compulsive disorder, obsessive-compulsive symptoms in the population, and insulin signaling"

**Supplementary Table 2.** Content of two novel OCS factors that could be compiled from the Toronto Obsessive-Compulsive Scale (TOCS) questions and that are similar in the questions they contain to the two factors (‘symmetry/counting/ordering’ and ‘contamination/cleaning’) from the Philadelphia Neurodevelopmental Cohort (PNC) data.

**Factor Symmetry/counting/ordering _TOCS_**

TOCS items:

“Do Certain”: Needs to do certain things (e.g., counting steps)

“Checks”: Checks things

“Count”: Needs to count objects

“Symmetry”: Needs things to be symmetrical

“Interfere”: Gets upset when people rearrange things

“Repeat”: Repeats actions until just right

“Not Exactly”: Worries if something done not exactly right

**Factor Contamination/cleaning _TOCS_**

TOCS items:

“Dirt”: Hates dirt/dirty things

“Germs”: Worries about germs

“Wash”: Needs to wash hands

“Clean”: Worries about being clean
