## Supplementary Table 3 for "Shared genetic etiology between obsessive-compulsive disorder, obsessive-compulsive symptoms in the population, and insulin signaling"

| **Supplementary Table 3.** Gene-wide association results for 51 unique genes from the OCD landscape and the OCS factors ‘symmetry/counting/ordering’ ^#^ | | | |
| --- | --- | --- | --- |
| Gene | P-value | Gene | P-value |
| *ADD3*; *ARHGAP15*; *BDNF*; *BTBD3*; *CCNC*; *CYTIP*; *DCC*; *DLGAP1*; *DLGAP3*; *DNAI1*; *DOCK1*; *EBF2*; *EFNA5*; *EREG*; *FKBP1A*; *GJD2*; *GNRH1*; *GRIN2B*; *HOXB8*; *HTR1B*; *IGF1*; *IGF1R*; *IRS2*; *ITGA9*; *KCNB2*; *KCNQ1* | 0.32005; 0.50966; 0.24211; 0.00680; 0.13363; 0.11929; 0.41032; 0.48661; 0.10051; 0.51632; 0.25946; 0.55613; 0.37218; 0.29452; 0.53514; 0.18184; 0.19447; 0.02840; 0.25101; 0.92934; 0.03339; 0.58297; 0.50513; 0.79192; 0.23801; 0.82367 | *LNX1*; *MEIS2*; *MTUS2*; *NOS1*; *NSG2*; *PDE4D*; *PRDM13*; *RACGAP1*; *REXO1*; *SEMA4D*; *SERPINH1*; *SLC1A1*; *SLIT3*; *SLITRK5*; *SORBS1*; *TBP*; *TFDP2*; *TMEM252*; *TNF*; *TRIOBP*; *TSPAN14*; *TXNL1*; *UBL3*; *ZBTB43*; *ZFP64* | 0.52292; 0.51374; 0.20290; 0.43244; 0.90462; 0.77885; 0.28219; 0.76819; 0.37348; 0.24246; 0.34515; 0.16768; 0.28663; 0.42304; 0.06008; 0.25962; 0.16867; 0.10289; 0.86862; 0.78470; 0.41173; 0.95027; 0.10305; 0.88721; 0.37194 |

^#^Note: Shown in this table are the gene-wide results from the MAGMA analyses of the 51 genes from the molecular landscape of OCD and the OCS factor ‘symmetry/counting/ordering’. None of the individual genes reached the Bonferroni-corrected P-value threshold of significance (P=0.05/51=0.00098).
