## Supplementary Table 4 for "Shared genetic etiology between obsessive-compulsive disorder, obsessive-compulsive symptoms in the population, and insulin signaling"

| **Supplementary Table 4**. PRS-based results for shared genetic etiology between OCD and two novel TOCS-based OCS factors and between five peripheral insulin signaling-related traits and two novel TOCS-based OCS factors. | | | | | |
| --- | --- | --- | --- | --- | --- |
| ‘base’ sample | ‘target’ sample | P_T_ | P-value | R^2^ | nSNPs |
| OCD | **Symmetry/counting/ordering _TOCS_** | **0.3** | **2,42E-05** | **0.49%** | **149119** |
|  | **Contamination/cleaning _TOCS_** | **0.4** | **4.07E-03** | **0.23%** | **185917** |
| Type 2 Diabetes | Symmetry/counting/ordering _TOCS_ | 0.1 | 3.39E-01 | 0.03% | 66870 |
|  | **Contamination/cleaning _TOCS_** | **0.1** | **1.59E-03** | **0.28%** | **67614** |
| HbA1c | Symmetry/counting/ordering _TOCS_ | 0.05 | 2.71E-01 | 0.04% | 20779 |
|  | Contamination/cleaning _TOCS_ | 0.001 | 1.68E-01 | 0.05% | 1018 |
| Fasting insulin | Symmetry/counting/ordering _TOCS_ | 0.05 | 3.71E-01 | 0.02% | 3926 |
|  | Contamination/cleaning _TOCS_ | 0.05 | 4.37E-01 | 0.01% | 3965 |
| Fasting glucose | Symmetry/counting/ordering _TOCS_ | 0.001 | 2.47E-01 | 0.04% | 484 |
|  | Contamination/cleaning _TOCS_ | 0.5 | 4.36E-01 | 0.01% | 21851 |
| 2 h Glucose | Symmetry/counting/ordering _TOCS_ | 0.05 | 4.45E-01 | 0.01% | 2924 |
|  | Contamination/cleaning _TOCS_ | 0.2 | 3.76E-01 | 0.02% | 9344 |

^#^ Shown in this table are the best SNP P-value thresholds (P_T_) for the PRSice analyses between OCD and two novel TOCS-based OCS factors and between five peripheral insulin signalling-related traits and two novel TOCS-based OCS factors, their Benjamini and Hochberg adjusted P-values for shared genetic etiology (P-value), the variance explained in the target sample phenotypes (R^2^), and the number of SNPs (nSNPs). Significant findings are indicated in bold. Abbreviations: TOCS=Toronto Obsessive-Compulsive Scale
