## Supplementary Figure 1A for "Shared genetic etiology between obsessive-compulsive disorder, obsessive-compulsive symptoms in the population, and insulin signaling"

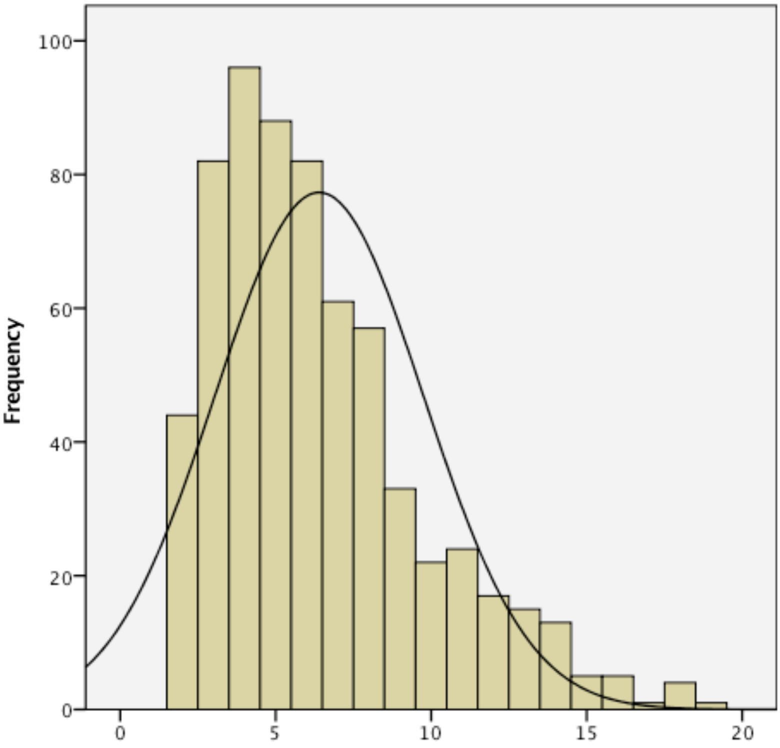


**Supplementary Figure 1A**. Histogram showing the distribution of the total OCS score in 650 children and adolescents aged 8-21 in the Philadelphia Neurodevelopmental Cohort (288 males and 362 females).
