## Supplementary Figure 1B for "Shared genetic etiology between obsessive-compulsive disorder, obsessive-compulsive symptoms in the population, and insulin signaling"

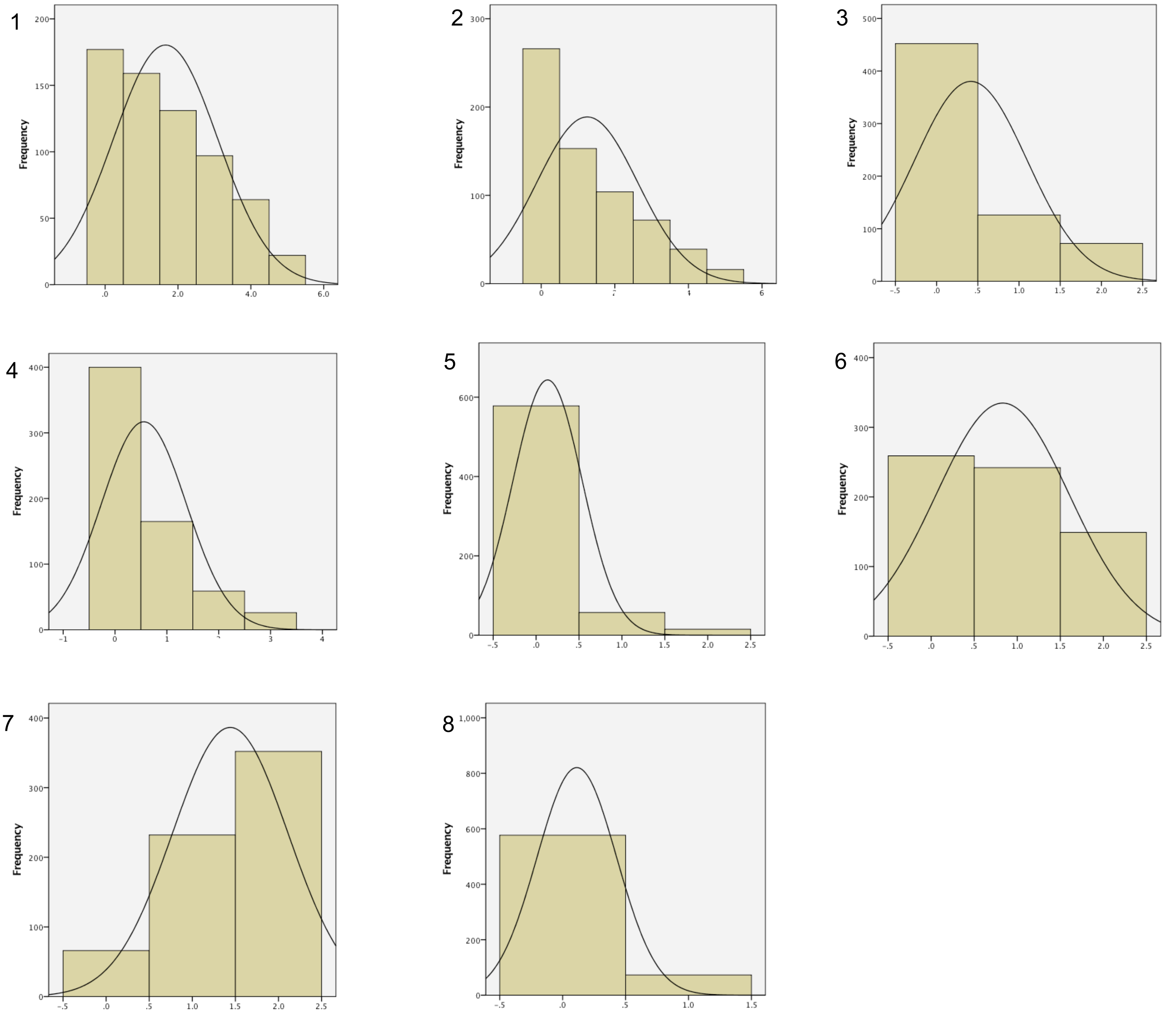


**Supplementary Figure 1B**. Histograms showing the distributions of the scores on eight OCS factors - that add up to the total OCS score - in 650 children and adolescents aged 8-21 in the Philadelphia Neurodevelopmental Cohort (288 males and 362 females): 1 'Impairment', 2 'Symmetry/counting/ordering', 3 'Contamination/cleaning', 4 'Aggressive taboo thoughts', 5 'Repetition', 6 'Guilty taboo thoughts, 7 'Distress', and 8 'Religious taboo thoughts'.
