## Supplementary Figure 3E for "Shared genetic etiology between obsessive-compulsive disorder, obsessive-compulsive symptoms in the population, and insulin signaling"

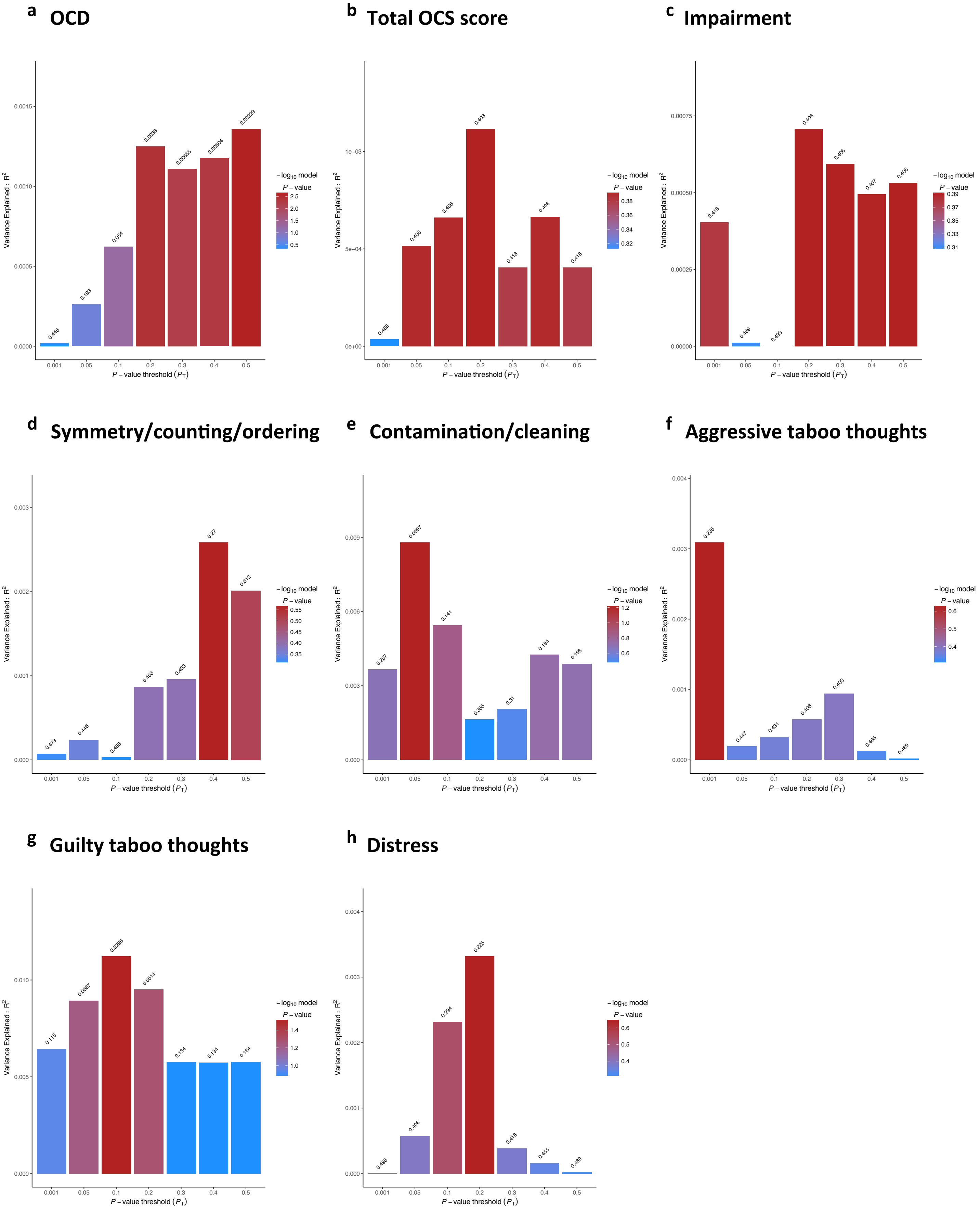


**Supplementary Figure 3E**. Bar plots from PRSice showing results at seven broad P-value thresholds ($P_T$) for shared genetic etiology between 2 h Glucose and obsessive-compulsive disorder (OCD), the total obsessive-compulsive symptom (OCS) score as well as six OCS factors (a–h) (see Methods). The numbers above the bars indicate the P-values for shared genetic etiology, and these P-values were corrected using the Benjamini-Hochberg false discovery rate method.
