## Supplementary Figure 4A for "Shared genetic etiology between obsessive-compulsive disorder, obsessive-compulsive symptoms in the population, and insulin signaling"

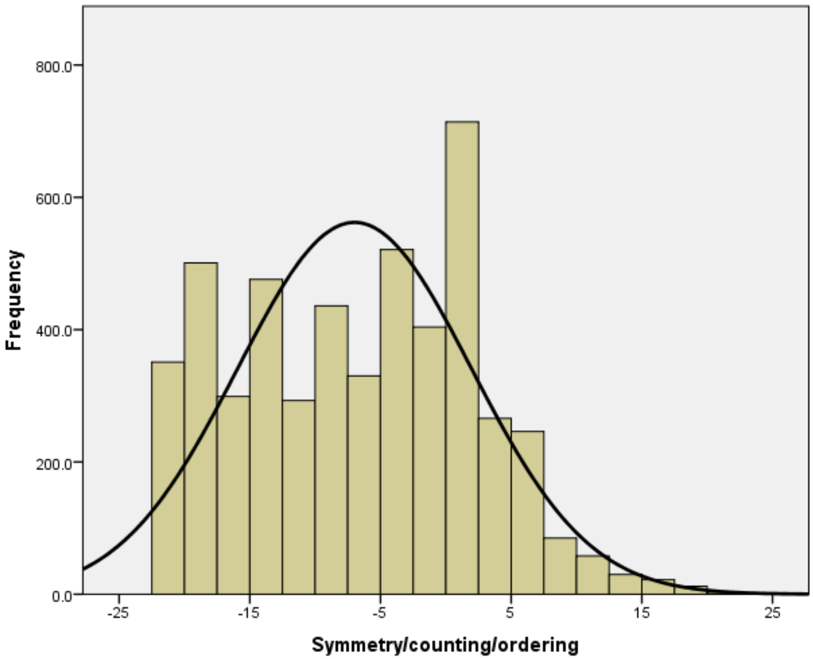


Supplementary Figure 4A. Histogram showing the distribution of the 'symmetry/counting/ordering' score in 5047 children and adolescents aged 6-17 in the Spit for Science Cohort.
