## Supplementary Figure 5C for "Shared genetic etiology between obsessive-compulsive disorder, obsessive-compulsive symptoms in the population, and insulin signaling"

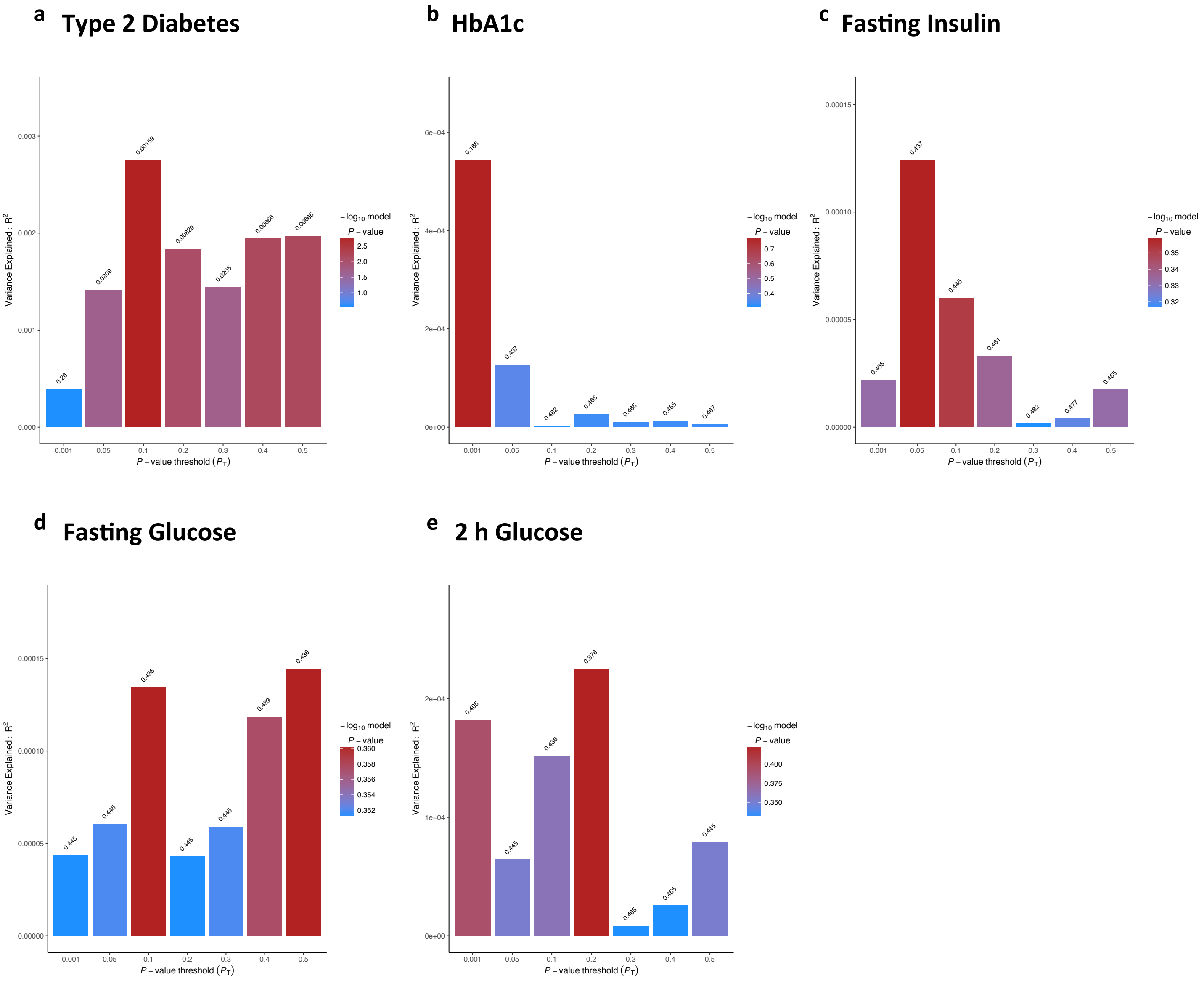


**Supplementary Figure 5C**. Bar plots from PRSice showing results at seven broad P-value thresholds ($P_T$) for shared genetic etiology between five peripheral insulin signaling-related traits (Type 2 Diabetes, HbA1C blood levels and blood levels of fasting insulin, fasting glucose and 2h Glucose) and TOCS factor ‘contamination/cleaning’ (a–e) (see Methods). The numbers above the bars indicate the P-values for shared genetic etiology, and these P-values were corrected using the Benjamini-Hochberg false discovery rate method.
